# Unified Protein-Small Molecule Graph Neural Networks for Binding Site Prediction

**DOI:** 10.1101/2025.09.03.674017

**Authors:** Jian Wang, Nikolay V. Dokholyan

## Abstract

Predicting small molecule binding sites on proteins remains a key challenge in structure-based drug discovery. While AlphaFold3 has transformed protein structure prediction, accurate identification of functional sites such as ligand binding pockets remains a distinct and unresolved problem. Graph neural networks have emerged as promising tools for this task, but most current approaches focus on local structural features and are trained on relatively small datasets, limiting their ability to model long-range protein–ligand interactions. Here, we develop YuelPocket, a new graph neural network that addresses these limitations through an innovative global graph design featuring a virtual joint node connecting all protein residues and small molecule atoms to capture long-range interactions while maintaining linear computational complexity. Trained on the comprehensive MOAD dataset containing over 38,000 protein-small molecule complexes, YuelPocket achieves exceptional performance with AUC-ROC values of 0.85 and 0.89 on COACH420 and Holo4k datasets, respectively.

## INTRODUCTION

Small molecule binding site prediction^1^ on proteins represents one of the most critical challenges in computational biology and drug discovery, with far-reaching implications for understanding protein function, rational drug design^2,3^, and structure-based drug discovery pipelines^4–7^. The accurate identification of binding pockets is fundamental to virtually every aspect of modern drug development, from target validation^8^ to lead optimization and clinical candidate selection.

The pharmaceutical industry faces unprecedented challenges in drug discovery, with development costs exceeding $2.6 billion per approved drug and failure rates nearing 90% in clinical trials^9^. A major factor contributing to these failures is the difficulty in accurately predicting protein-small molecule interactions. Traditional drug discovery methods often depend heavily on experimental techniques, which are costly and time-consuming. The introduction of AlphaFold2^10^ has transformed protein structure prediction, achieving unmatched accuracy in determining three-dimensional protein structures from amino acid sequences. The release of AlphaFold3^11^ has further enabled researchers to directly predict protein-small molecule complexes, potentially significantly speeding up drug discovery. However, despite its impressive capabilities, AlphaFold3 still encounters notable challenges in drug discovery. Shen et al.^12^ found that while AlphaFold3 reliably predicted the overall structure of receptors, its precision in positioning small molecule ligands was variable and frequently inaccurate. This inconsistency was especially evident with allosteric modulators, highlighting a key obstacle in reliably identifying binding sites.

Existing pocket prediction algorithms can be broadly categorized into three main approaches, each with significant limitations. Traditional geometric approaches such as Fpocket^1^, SiteHound^13^, and CASTp^14^ rely on geometric descriptors including surface curvature, solvent accessibility, and cavity detection algorithms. These methods identify potential binding sites based on physical properties such as surface concavity, pocket depth, solvent accessibility, and geometric complementarity to spherical probes. While these methods offer fast computation and interpretable results without requiring training data, they suffer from limited accuracy and poor performance on flat binding sites or allosteric sites.

More recent machine learning approaches employ various techniques to improve binding site prediction accuracy. P2Rank^15^ uses a Random Forest classifier with geometric and evolutionary features to predict ligand binding sites on a protein’s solvent-accessible surface, trained on the CHEN11^16^ dataset containing 251 proteins with 476 ligands. DeepSite^17^ utilizes 3D convolutional neural networks that process 16 Å^3^ protein subgrids with eight feature channels, trained on the scPDB v.2013 database with 7,622 binding sites. DeepPocket^18^ employs 3D Convolutional Neural Networks for binding site detection and segmentation, trained on the scPDB^19^ v.2017 database with 17,594 binding sites from 16,612 proteins. PUResNet^20^ combines U-Net^21^ and ResNet^22^ architectures to predict binding site probabilities for each voxel in 3D protein structures. While these methods offer improved accuracy over geometric approaches, they are limited by computational intensity and difficulty in capturing long-range interactions.

Emerging graph neural network (GNN)^23^-based approaches provide more sophisticated modeling of protein structures, but suffer from fundamental limitations in both architecture and training data scale. PocketMiner^24^ uses a geometric vector perceptron graph neural network to predict cryptic pockets, though it was trained on only 37 proteins from molecular dynamics simulations, severely limiting its generalization capabilities. SiteRadar^25^ represents protein structures as graphs where nodes are heavy atoms and edges are interatomic distances, predicting binding sites on a grid by classifying each grid point as pocket or non-pocket. LigBind^26^ employs a relation-aware graph neural network to predict ligand-specific binding residues, representing each residue and its surrounding structural context (within 15 Å radius) as a graph, with a two-phase transfer learning approach trained on over 1,000 ligands. While these methods represent advances over traditional approaches, they share critical architectural limitations: they construct only local graphs that capture limited spatial relationships between neighboring elements, fundamentally missing the global interaction patterns and synergistic binding effects where distant residues cooperatively contribute to ligand recognition. Particularly, SiteRadar creates graphs between grid points and their surrounding protein atoms, while LigBind builds graphs between residues and their neighboring protein residues within a fixed radius. Furthermore, these methods do not explicitly incorporate ligand information during the prediction process, limiting their ability to predict ligand-specific binding sites, and their training datasets remain relatively small compared to the diversity of protein-small molecule interactions in nature.

Here, we develop an innovative graph neural network architecture, YuelPocket, to address these fundamental limitations through a global protein-small molecule interaction graph (Figure 1). Unlike existing GNN approaches that rely on local graph construction, we introduce a virtual joint node (Figure 1c) that serves as a global information hub, connecting all protein residues and all ligand atoms in a unified computational framework. The virtual joint node acts as a computational bridge that aggregates information from all protein residues and ligand atoms, and distributes the processed information back to the protein and the compounds. This design enables the model to capture the synergistic nature of molecular recognition while maintaining linear computational complexity *O(C+m+n)* instead of the quadratic complexity *O(C+m×n)* that would result from direct all-to-all connections between protein residues and compound atoms, where *C* represents the summation of the number of protein backbones (Figure 1a), residue contacts (Figure 1a), and compound bonds (Figure 1b), *m* represents the number of protein residues, and *n* represents the number of ligand atoms.

**Figure 1.**
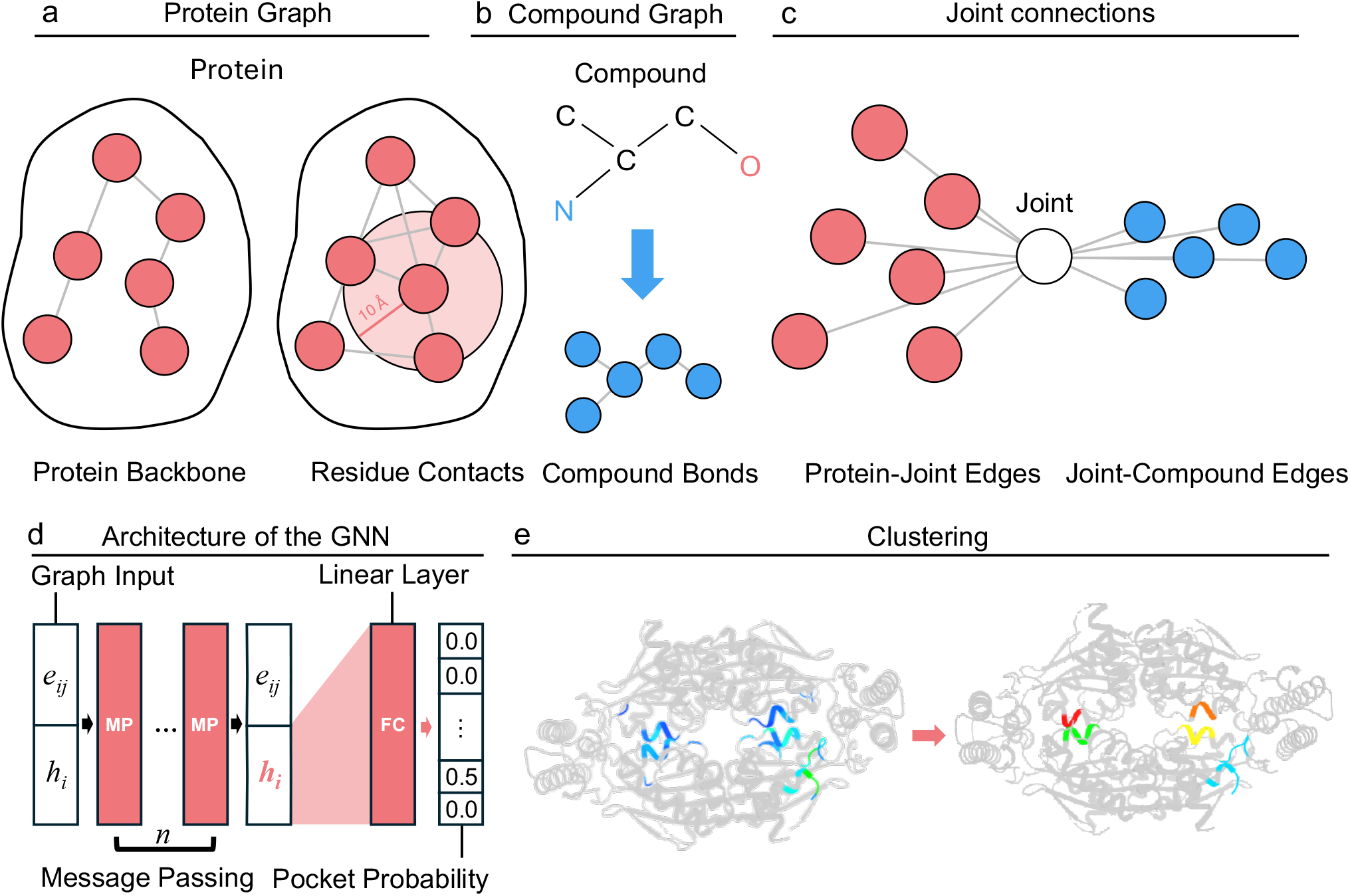
Structural and topological representation of protein-compound interactions in YuelPocket’s graph neural network framework. (a) Protein graph: Red nodes depict Cα atoms (spheres) with edges (black lines) representing backbone connectivity (labeled “Backbone”) and inter-residue contacts (“Contacts”). (b) Compound graph: Blue nodes show heavy atoms (C, N labeled) with bond edges (“Bonds”); stick model overlay illustrates chemical structure. (c) Joint connection topology: Virtual joint node (labeled “Joint”) mediates protein-compound interactions via two edge types: “Protein-Joint Edges” and “Joint-Compound Edges”, avoiding all-to-all residue-atom connections. (d) GNN architecture: Graph input (left) passes through message-passing layers (center) with learned edge weight updates, culminating in pocket probability predictions (right). (e) Spatial clustering: Euclidean embedding of protein residues (colored by cluster ID) reveals pocket localization patterns.

We tested YuelPocket on two commonly used datasets for small molecule binding site prediction evaluation, COACH420^15^ and Holo4k^15^. Our model achieves AUC-ROC values of 0.85 and 0.89, respectively, on these datasets, significantly outperforming random guessing (0.5) and demonstrating robust discrimination between pocket and non-pocket residues. We observe exceptional specificity (0.85) and NPV (0.95), indicating the ability of YuelPocket to accurately identify non-binding regions while minimizing false positives. These results suggest that our global graph architecture successfully captures the complex patterns underlying protein-small molecule interactions, enabling precise binding site prediction. Further, the ligand-specific predictions of YuelPocket, which we demonstrate through visualization of different ligands binding to the same protein, confirm its ability to capture the unique characteristics of individual protein-small molecule interactions rather than predicting generic binding sites. We also developed a minimal probe set approach using 15 selected ligands to comprehensively explore binding space across diverse protein targets, implying the potential for high-throughput drug discovery applications.

## Results

### Binding Site Prediction Performance on Benchmark Datasets

We conducted the training and evaluation using the comprehensive Binding MOAD^27^ dataset, which contains over 38,000 protein-small molecule complexes from the Protein Data Bank^19^. For performance assessment, we focused on two widely-used benchmark datasets: COACH420 and Holo4k, the proteins in which are also covered by MOAD. We split the MOAD dataset into training and testing sets, using proteins from COACH420 and Holo4k as the test set and the remaining proteins as the training set, resulting in a training-to-testing ratio of approximately 8.5:1.5. We define pocket residues as those within 6 Å of any ligand atom. YuelPocket outputs a probability score (0-1) for each residue being part of the binding pocket.

We evaluated YuelPocket’s performance using ROC (Receiver Operating Characteristic) curves (Figure 2a) and precision-recall curves (Figure 2b) on COACH420 and Holo4k datasets. YuelPocket achieves AUC-ROC (area under the ROC curve) values of 0.85 and 0.89 on the two datasets, respectively, significantly outperforming random guessing (0.5). The AUC-PR values are 0.49 and 0.46, respectively, which are much lower than the AUC-ROC due to the severe class imbalance in the dataset, where pocket residues represent only a small fraction of total residues. We computed the AUC-PR for random guessing as 0.053, confirming the imbalanced nature of the datasets. Despite this challenge, the AUC-PR values are substantially higher than random guessing, demonstrating YuelPocket’s ability to identify binding pocket residues while maintaining high precision.

**Figure 2.**
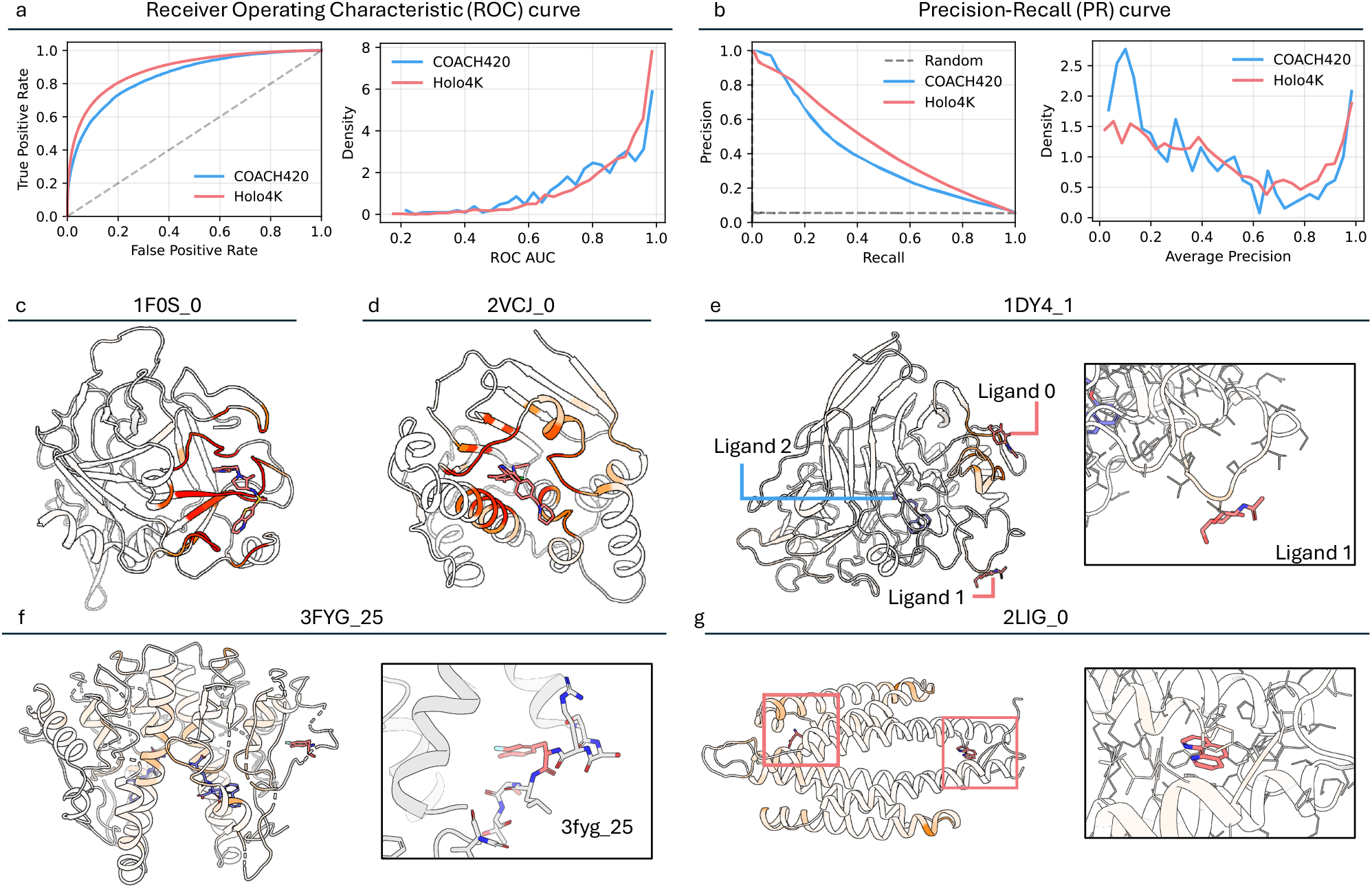
Performance evaluation and binding site prediction visualization of YuelPocket. (a) Receiver Operating Characteristic (ROC) curves showing AUC-ROC values of 0.85 (COACH420) and 0.89 (Holo4k), significantly outperforming random guessing (dashed line at 0.5). (b) Precision-Recall (PR) curves with AUC-PR values of 0.49 (COACH420) and 0.48 (Holo4k), demonstrating robust performance despite severe class imbalance (random guessing AUC-PR = 0.05). (c-g) PyMOL visualizations of five representative protein-ligand complexes from MOAD, with residues colored by predicted binding probability (red: high probability; white: low probability). Protein backbones are shown as gray cartoons, with ligand molecules as sticks. Ligand-specific prediction of 1DY4_1 (e) showing correct identification for ligand 1 but not ligand 2. Case 3FYG_25 highlights MOAD annotation error (ligand is actually a modified residue).

To visualize our predictions, we color residues based on their predicted probability in PyMOL^28^. For example, in proteins 1F0S (Figure 2c) and 2VCJ (Figure 2d), residues around ligand atoms are predominantly colored red (close to 1), indicating correct prediction of pocket residues. Importantly, our predictions are ligand-specific, meaning pocket residues vary for different ligands. For instance, protein 1DY4 has three ligands: 1DY4_0 and 1DY4_1 are identical ligands, while 1DY4_2 is a different ligand molecule. In the MOAD dataset, each protein-ligand complex is identified by a unique code following the format “PDBID_LigandIndex”, where PDBID is the four-character Protein Data Bank identifier and LigandIndex indicates the specific ligand molecule (0, 1, 2, …) bound to that protein. When we use 1DY4_1 as the input ligand, residues around 1DY4_0 appear mostly red, residues around 1DY4_1 show slight red coloring, while residues around 1DY4_2 remain white, demonstrating ligand-specific prediction. Residues around 1DY4_0 and 1DY4_1, both showing red coloring, indicate successful prediction for the same ligand type. The subtle red coloring for 1DY4_1 is probably because this binding site is not a deep cavity but rather a shallow binding site with minimal residue-ligand interactions. This observation raises an interesting possibility: the pocket probability may correlate with binding affinity, as shallow binding sites typically exhibit lower binding affinities than deep cavities. We investigate this relationship in detail in the following section.

The low AUC-PR values primarily result from the severe class imbalance in the dataset. We also identified two additional factors contributing to this limitation. First, we discovered inherent errors in the MOAD dataset. For example, the ligand 3FYG_25 (Figure 2f) is actually a modified residue in the original PDB file, but the MOAD dataset incorrectly extracts this residue and treats it as a ligand that is not covalently bonded with the protein. Second, many proteins likely contain unknown binding pockets that have not been experimentally characterized. When YuelPocket predicts these uncharacterized binding sites, they are counted as false positives. For instance, YuelPocket predicts multiple potential binding pockets for protein 2LIG_0, of which only two are experimentally validated, while the others represent potentially real but uncharacterized binding sites.

### Binary Classification Threshold Optimization

YuelPocket outputs probability scores (0-1) for each residue being part of the binding pocket. While these continuous probabilities provide detailed information about binding likelihood, many practical applications require binary pocket/non-pocket classifications for decision-making in drug discovery pipelines. For example, researchers need to know which specific residues to target for mutagenesis studies^29^, which regions to focus on during molecular docking^4,5^, or which areas to prioritize for experimental validation. We therefore convert these probabilities into binary classifications using probability thresholds. We first analyzed the distribution of predicted probabilities across all residues in the training set (Figure 3a) and identified outliers as potential pocket residues. This approach is based on the observation that binding pocket residues represent a small, distinct subset of protein residues with unique structural and chemical properties that enable ligand binding. In the probability distribution, these residues naturally form outliers because they exhibit significantly higher binding probabilities compared to the majority of non-binding residues. Since most outliers occur above 0.08 (Figure 3a), we established a fixed threshold of 0.08 for all proteins as our first strategy. However, we observed that probability distributions vary significantly across different proteins. For example, predicted probabilities for 1F0S_0 (Figure 3b) are predominantly below 0.05, while those for 1HI5_0 are mostly below 0.78. This variation necessitated a protein-specific approach. We therefore developed an adaptive-threshold strategy that sets the threshold for each protein based on its individual probability distribution, specifically using 1.5 times the interquartile range (IQR) above the median.

**Figure 3.**
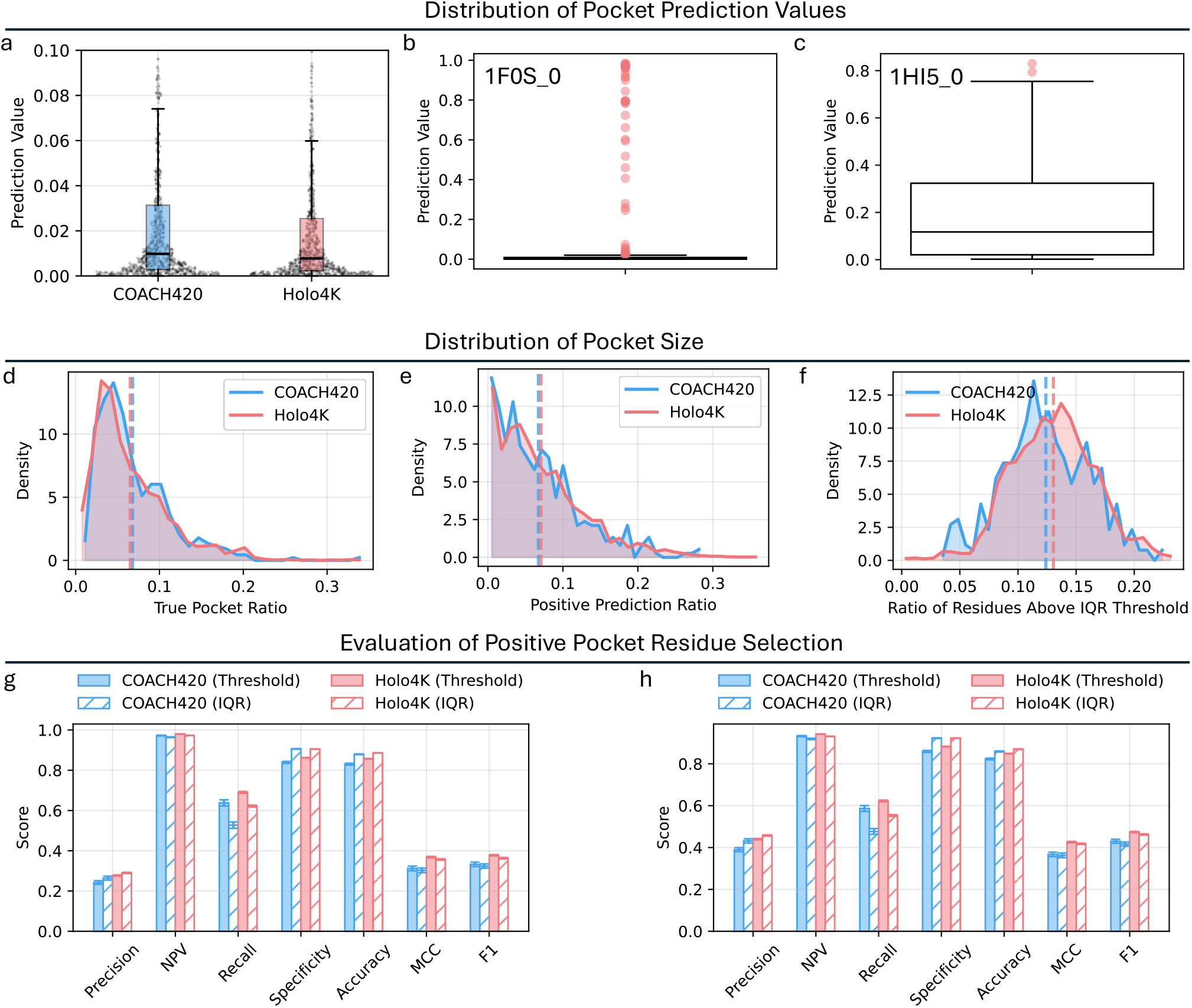
Threshold strategy analysis for binary classification of binding pockets in YuelPocket. (a) Histogram of predicted binding probabilities for all training set proteins. (b) 1F0S_0 exhibits low probabilities (90% <0.05). (c) 1HI5_0 shows broader distribution with outlier threshold at 0.78. (d-f) Ratios of true pocket residues (d), ratios of pocket residue predicted with fixed-threshold strategy (e), and ratios of pocket residues predicted with adaptive-threshold strategy (f) across proteins. (g) Precision, NPV, recall, specificity, accuracy, MCC, and F1 score of pocket residue prediction by using the two threshold selection strategies on the COACH420 and Holo4K datasets. Threshold means the fixed-threshold strategy. IQR means the adaptive-threshold strategy.

We first evaluated the ratio of predicted pocket residues with the two strategies. As a comparison, we calculated the true pocket residue ratio (residues within 6 Å of any ligand atom) across the training set (Figure 3d), finding that most proteins have ratios in the range of 0-0.2, with a median around 0.06. Both strategies produced predicted ratios that generally matched this distribution. However, the fixed-threshold strategy resulted in a portion of proteins with a near-zero number of pocket residues (Figure 3e), whereas the adaptive-threshold strategy and true pocket residues showed fewer such cases (Figure 3f). This difference occurs because the fixed threshold of 0.08 is too high for some proteins, preventing any residues from being classified as pocket residues.

Using these threshold strategies, we converted continuous probabilities into binary classifications and calculated standard performance metrics on the test set (Figure 3g). Our model achieved a negative predictive value (NPV) of 0.95 and a specificity of 0.85, demonstrating strong performance in identifying non-binding regions. Recall reached 0.6, indicating that our model captures a substantial portion of true binding sites. However, precision was around 0.25, with correspondingly low Matthews correlation coefficient (MCC) and F1-scores around 0.3. These results reflect the inherent challenges of binding site prediction, where the severe class imbalance (binding site residues represent only 3-15% of total residues) and the presence of uncharacterized binding sites contribute to lower precision scores. When our model predicts potentially real but uncharacterized binding sites, they are counted as false positives, artificially reducing precision metrics. Notably, when we include all pocket residues for all ligands of each protein, precision improves significantly to around 0.4, further supporting the existence of many uncharacterized binding pockets.

Comparing the two threshold strategies, we found that adaptive-threshold achieves higher precision, specificity, and accuracy than fixed-threshold while showing lower values for other metrics. This performance difference reflects the adaptive strategy’s ability to account for protein-specific characteristics and binding site diversity. The adaptive approach adjusts classification thresholds based on each protein’s individual probability distribution, better handling proteins with unusual binding site properties or multiple binding pockets of varying strengths. This protein-specific adaptation produces more conservative predictions, resulting in higher precision and specificity by reducing false positives. However, this conservative approach reduces recall and negative predictive value (NPV) by making the model more selective. For instance, when the protein is small (Figure S1), most residues are indeed pocket residues, yet the adaptive threshold predicts only a small subset of them (Figure S2), resulting in low recall. This precision-recall trade-off emphasizes the importance of selecting the appropriate threshold strategy based on application requirements.

### Binding Site Clustering and Center Prediction

While our previous results demonstrate that YuelPocket accurately predicts individual residues within 6Å of ligand atoms, researchers typically need to understand the overall location and number of binding pockets rather than specific residue-level predictions. Although visualization by coloring residues with predicted probabilities provides qualitative insights, we implemented a quantitative clustering approach to group predicted pocket residues into distinct binding sites. We applied hierarchical clustering to predict pocket residues based on their spatial distances. This approach groups spatially proximal residues that likely belong to the same binding site, providing quantitative insights into the number and location of potential binding pockets. We define a pocket as successfully predicted if the minimal distance between residues in that pocket and ligand atoms falls below a specific threshold. By varying the threshold from 0 to 20 Å, we calculated prediction success rates and found that the success rate rapidly increases to 90 % as the threshold increases from 0 to 3 Å (Figure 4a). This demonstrates YuelPocket’s high accuracy in predicting binding pocket locations. For example, in protein 1QJW (Figure 4b), all predicted pocket residues cluster into two distinct binding pockets that precisely align with its two ligand binding sites.

**Figure 4.**
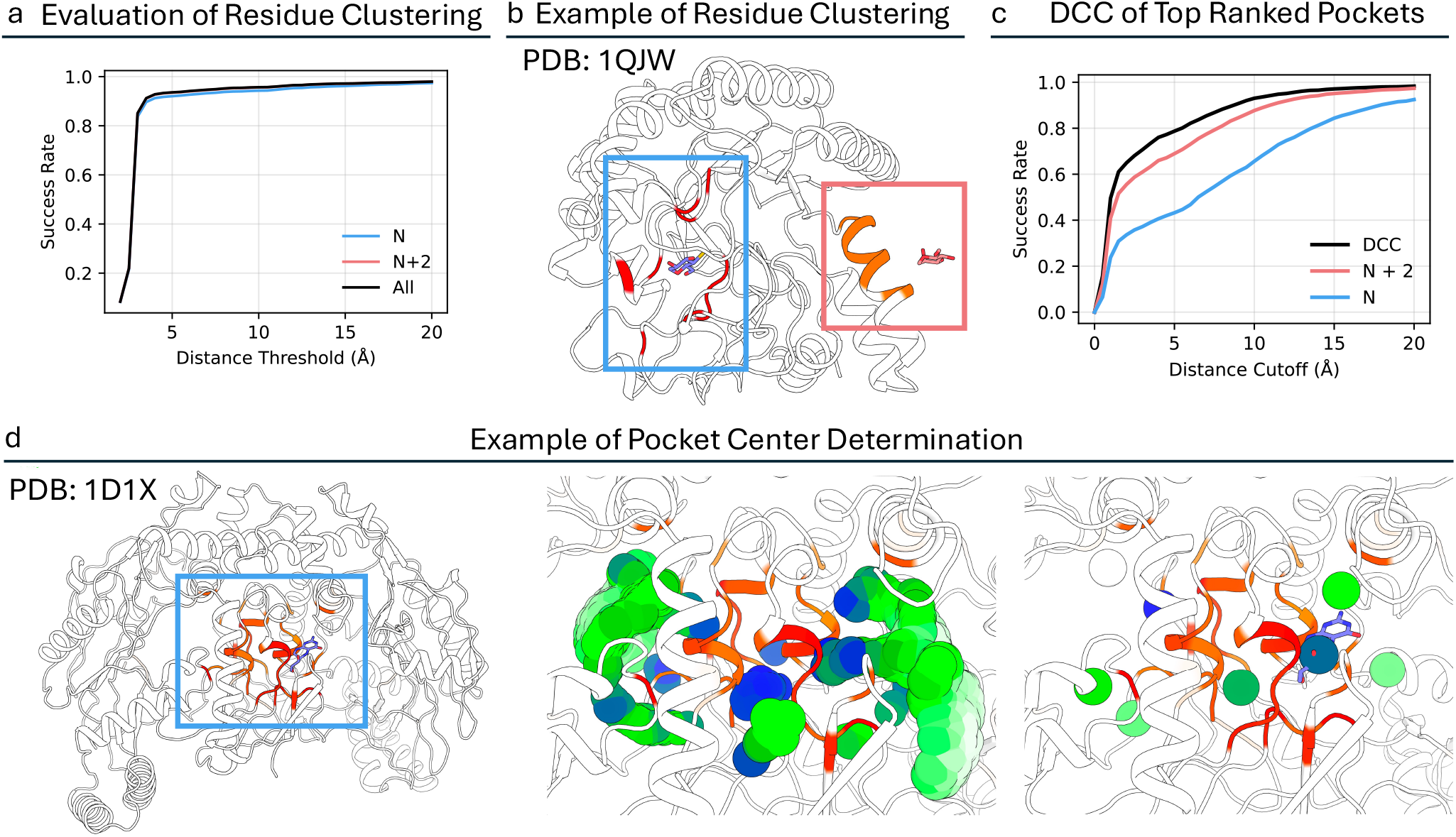
Binding pocket clustering analysis of YuelPocket predictions. (a) Success rates (y-axis) of top-ranked (Top N, Top N, and All) predicted pockets at varying distance cutoff threshold from true ligand atoms, where N is the number of ligands for a specific protein. (b) Clustering of the pocket residues of protein 1QJW. The blue box and red box show the two clusters. (c) Success rates (y-axis) of top-ranked (Top N, Top N, and All) predicted pockets with their centres at varying distance cutoff threshold from the center of the ligands. (d) Illustration of the pocket center determination algorithm. The spheres represent the clustering centroids of all grid points on the pocket surface (see Methods). The color of each sphere corresponds to the average pocket probability of the residues near it.

Most existing pocket prediction methods predict pocket center positions rather than residue-level binding sites. To enable direct comparison with these methods, we developed an algorithm (Methods) that predicts pocket centers based on our residue-level predictions. This algorithm creates a 3D scoring grid around the protein using pocket prediction scores and atom positions, then employs Fast Fourier Transform (FFT) convolution^30^ and DBSCAN^31^ clustering to identify potential binding site centers. This approach conceptually mirrors rigid docking algorithms, where ligand molecules are systematically positioned and scored across protein surfaces to find optimal binding locations by using FFT.

We evaluate prediction success by calculating the distance between predicted pocket centers and ligand centers (DCC), considering predictions successful if this distance is less than 5 Å. Since some pockets may be large and require multiple centers for adequate representation, we analyzed success rates under different conditions. When considering only the top-ranked pocket center (N), we achieved a 43 % success rate at the 5 Å threshold. Including the top N+2 ranked pocket centers increased the success rate to 69 %, while considering all predicted pocket centers yielded approximately 79 % success rate (Figure 4c, Table S1 & S2). To benchmark our performance against existing methods, we compared YuelPocket with representative algorithms from the three main categories of pocket prediction approaches (Table S2): Fpocket (geometric methods), SiteRadar (graph neural network methods), and PUResNet (other machine learning methods). Our comparison reveals that YuelPocket achieves competitive performance with a DCC score of 0.787, outperforming Fpocket (0.710), PUResNet (0.460), and SiteRadar (0.760).

Clustering and predicting pocket centers introduce additional algorithmic complexity and potential errors that are not inherent to our core prediction method. The accuracy of YuelPocket is primarily demonstrated through the ROC and PR curve analyses in the first section, while clustering and predicting pocket centers serve as a supplementary tool for structural analysis and method comparison.

### Comprehensive Binding Site Discovery with Minimal Probe Sets

For some proteins, it is challenging to develop drugs targeting the known active site due to structural constraints, functional requirements, or drug resistance mechanisms. In such cases, identifying allosteric sites^32^ or novel binding pockets becomes crucial for drug discovery. Since YuelPocket predicts ligand-specific binding sites, we developed a minimal probe set approach (Methods, Figure 5a) to comprehensively identify all potential binding sites across diverse protein targets. A minimal probe set represents an optimized collection of ligands that can collectively cover the majority of binding sites across the entire protein dataset while minimizing redundancy and computational cost.

**Figure 5.**
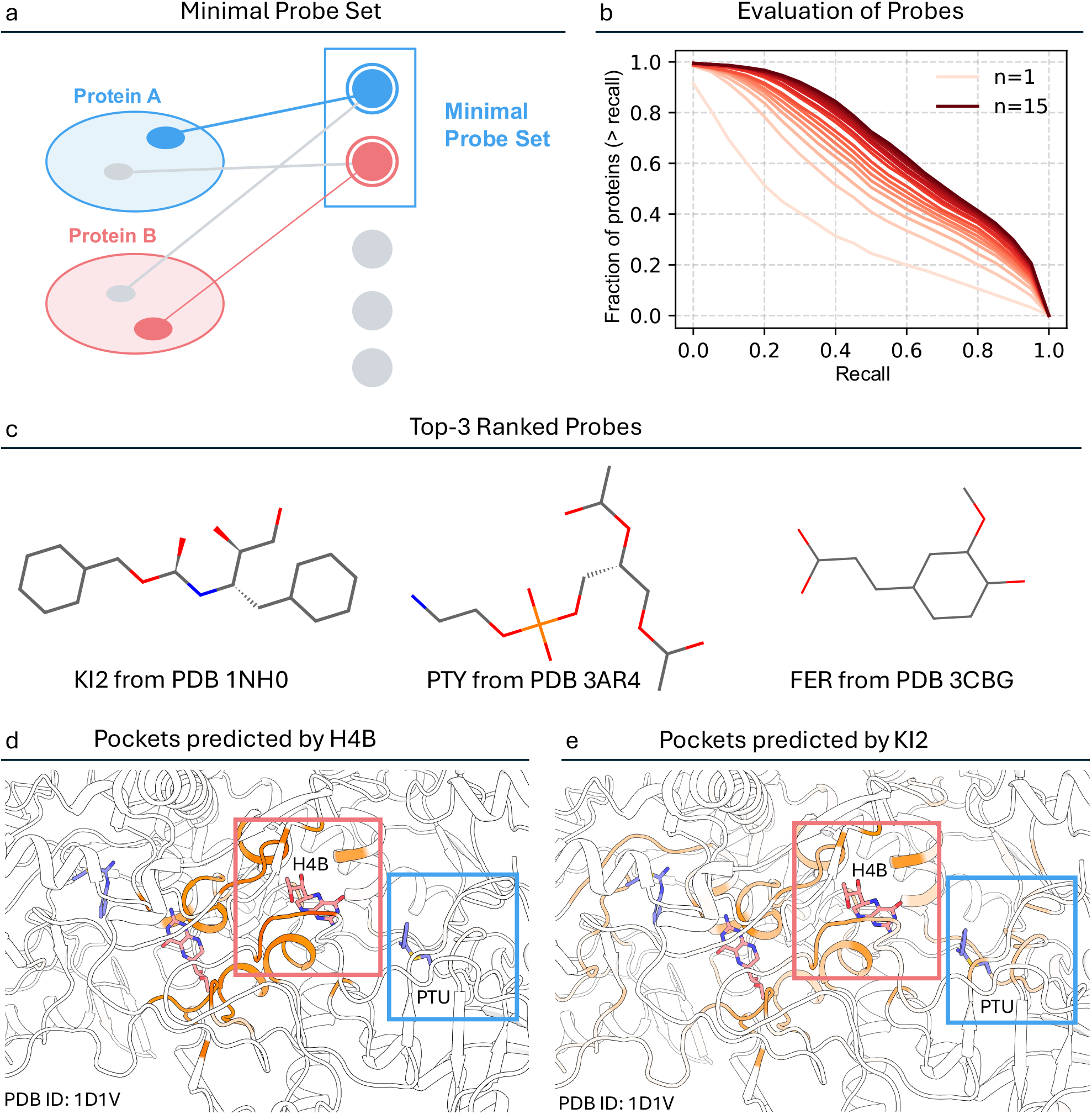
Minimal probe set construction and evaluation for comprehensive binding site mapping. (a) Schematic of the definition of minimal probe set. (b) Fraction of proteins with the recall of pocket residues greater than the recall threshold versus the recall threshold. The 15-probe set (red curve) achieves superior coverage compared to single probes. (c) Chemical structures of top 3 ranked probes: KI2 (from PDB 1NH0), PTY (from PDB 3AR4), and FER (from PDB 3CBG). (d) Native ligand H4B only detects its own binding site, missing the PTU pocket. (e) Probe KI2 identifies both H4B and PTU sites, demonstrating pan-pocket coverage.

We implemented a greedy algorithm to construct the minimal probe set, which operates by iteratively selecting ligands that provide the maximum incremental coverage of true pocket residues across the protein dataset. The algorithm begins with an empty probe set and systematically evaluates each candidate ligand based on its ability to identify previously uncovered binding site residues. At each iteration, the ligand that contributes the most new coverage is added to the set, and the process continues until a predefined coverage threshold is achieved. This approach resulted in a minimal probe set of 15 ligands (Top 3 shown in Figure 5c; All shown in Table S3) that provides comprehensive coverage of binding sites across diverse protein families. As the number of probes increases, a larger fraction of proteins achieve recall values above a given threshold for pocket residue identification, demonstrating the effectiveness of the probe set in capturing diverse binding sites (Figure 5b).

Compounds in the minimal probe set usually achieve higher coverage of pocket residues than the other compounds. For example, in protein 1D1V, the known ligand H4B primarily identifies residues around its own binding site, missing the pocket for ligand PTU entirely (Figure 5d). However, when using probe KI2 from our minimal set, the model successfully identifies both the H4B and PTU binding sites, demonstrating the probe’s ability to capture multiple binding pockets within a single protein (Figure 5e). While the prediction probabilities for the H4B site decrease slightly when using KI2, this trade-off is acceptable given the significant gain in overall binding site coverage.

### Correlation between Pocket Probability and Binding Affinity

We also explored whether the probability of finding a binding pocket for a specific ligand on a protein correlates with the binding affinity between that ligand and protein. This question is particularly significant because YuelPocket was trained exclusively on structural information (protein 3D coordinates and ligand 2D structures) without any exposure to binding affinity data during training. We investigated this relationship using the PDBBind dataset, which contains 5,314 protein-ligand complex structures along with their corresponding Kd and Ki values. For each complex, we computed pocket probabilities using YuelPocket. Since YuelPocket outputs probability scores for individual residues, we employed two aggregation methods to assess correlation with binding affinity: selecting the maximum probability across all residues and calculating the mean probability across all residues.

Our analysis revealed significant correlations between pocket prediction probabilities and binding affinities, despite the model never having been trained on affinity data (Figure S3). For maximum probability analysis, we observed Pearson correlation of 0.391 and Spearman correlation of 0.423 across all samples (Table S4). Mean probability analysis yielded even stronger correlations, with a Pearson correlation of 0.429 and a Spearman correlation of 0.415 across all samples. These results demonstrate that YuelPocket’s pocket prediction probabilities exhibit moderate to strong correlations with experimental binding affinities, particularly when using mean probability aggregation. This correlation is remarkable because it emerges purely from structural learning without any explicit affinity information during training.

The emergence of binding affinity correlations from purely structural training data has profound implications for our understanding of protein-ligand interactions. It suggests that the structural features that determine binding site formation are inherently linked to the energetic factors that govern binding strength. This finding further corroborates the fact that the interaction energy must overcome a significant entropic loss due to the binding of a small molecule. Our virtual joint node architecture, by learning to represent the binding pocket as a distinct molecular environment, appears to capture these underlying physical principles that connect structure to function.

## Discussion

The introduction of the virtual joint node addresses the computational complexity challenge in protein-ligand interaction modeling. An intuitive approach to construct a unified graph is to directly connect all protein residues to all ligand atoms, but it will create a quadratic scaling problem that becomes computationally intractable for large proteins or complex ligands. The edge count in such approach is C + m×n, where *C* represents the summation of the number of protein backbones (Figure 1a), residue contacts (Figure 1a), and compound bonds (Figure 1b), m represents the number of protein residues, and n represents the number of ligand atoms. The m×n term creates quadratic complexity that grows rapidly with molecular size, limiting the applicability of these methods to small proteins and simple ligands. We introduce the virtual joint node as an intermediary that dramatically reduces computational complexity while maintaining information flow between protein and ligand components. In YuelPocket, the edge count is C + m + n, resulting in linear complexity that scales efficiently with molecular size. This reduction enables the model to handle large proteins and complex ligands while maintaining the ability to capture the essential interactions that define binding specificity.

Our results show that the same protein can exhibit dramatically different binding site predictions depending on the input ligand, as demonstrated by the 1DY4 example, where 1DY4_1 and 1DY4_2 produce distinct binding site patterns. The subtle red coloring observed for 1DY4_1, despite being identical to 1DY4_0, highlights the model’s sensitivity to binding site characteristics: shallow binding sites with minimal residue-ligand interactions produce lower prediction probabilities, reflecting the reduced binding affinity typically associated with such sites.

The discovery of annotation errors in the MOAD dataset, such as the 3FYG_25 case, where a modified residue was incorrectly classified as a ligand, raises important questions about dataset quality in computational biology. These errors artificially inflate false positive rates and contribute to the observed precision-recall imbalance. More significantly, our analysis suggests that many proteins contain uncharacterized binding sites that are counted as false positives when predicted by our model. The precision improvement from 0.2 to 0.4 when considering all ligands for a protein supports this hypothesis, indicating that our model may be identifying real but experimentally unvalidated binding sites.

The comparison between fixed-threshold and adaptive-threshold strategies reveals important trade-offs that must be considered based on specific application requirements. The adaptive-threshold approach’s superior precision and specificity make it ideal for high-confidence applications where false positives are costly, such as target validation. However, the reduced recall associated with this conservative approach may be problematic for comprehensive screening applications where missing potential binding sites is more concerning than false positives. The protein-specific nature of probability distributions, exemplified by the dramatic differences between 1F0S_0 (mostly <0.05) and 1HI5_0 (mostly <0.78), suggests that adaptive thresholds are essential for proteins with unusual binding characteristics or multiple binding pockets of varying strengths.

The clustering analysis reveals fundamental differences between residue-level prediction methods like YuelPocket and traditional center-based approaches. While our success rate in pocket location prediction demonstrates excellent performance, the lower success rates in center prediction highlight the challenges of comparing fundamentally different prediction paradigms. The success rate improvement when considering all predicted centers suggests that our method identifies multiple potential binding sites within large pockets, which may represent distinct sub-pockets or alternative binding modes. This granularity provides valuable information for drug design but complicates direct comparison with methods that predict single pocket centers.

In conclusion, YuelPocket demonstrates effective small molecule binding site prediction with AUC-ROC values of 0.85-0.89 on benchmark datasets. The method shows ligand-specific prediction capabilities and reveals correlations between pocket probabilities and binding affinities despite being trained only on structural data. The adaptive threshold strategy provides better precision than fixed thresholds, while clustering analysis enables comparison with traditional pocket prediction methods.

## Methods

### Raw Data Collection

We employ a multi-stage approach to transform three-dimensional structural information into graph representations suitable for deep learning. For each protein-ligand complex, we extract atomic coordinates, residue types, and chemical connectivity patterns through specialized parsing functions.

PDB files are processed to extract residue-level information, where each residue is represented by its Cα atom coordinates x_*i*_ ∈ ℝ^3^ and one-hot encoded amino acid type 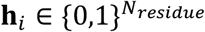. The backbone connectivity is established through spatial proximity analysis, where consecutive residues *i* and *j* are connected if their Cα atoms satisfy the distance constraint ||**x**_*i*_ − **x**_*j*_||_2_ < 4.1 Å.

### Pocket Detection and Ground Truth Generation

We employ a distance-based approach to identify protein residues that form the binding interface with the ligand. For each protein residue *i*, we calculate the minimum distance to any ligand atom:

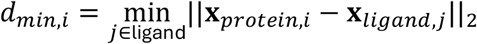

A residue is classified as part of the binding pocket if *dm*_*in,i*_ ≤ *τ*_*pocket*_, where *τ*_*pocket*_ = 6.0 Å represents the interaction threshold. This generates binary pocket labels 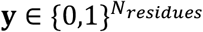 that serve as ground truth for supervised learning:

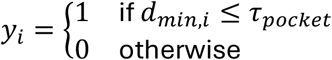

### Graph Construction

The YuelPocket graph neural network operates on a heterogeneous graph that represents both protein and ligand structures in a unified computational framework. The graph construction process begins with the extraction of protein three-dimensional coordinates from PDB files and ligand two-dimensional structural information from SMILES^33^ strings or molecular files. Each protein residue is represented as a node in the protein subgraph, with node features encoding amino acid type, secondary structure, solvent accessibility, and spatial coordinates. Edges between protein residues are established based on spatial proximity, typically connecting residues within 8 Å of each other, capturing both covalent and non-covalent interactions that define the protein’s structural context.

The ligand subgraph represents each atom as a node with features including atom type, hybridization state, formal charge, and aromaticity. Edges connect bonded atoms, preserving the molecular connectivity and electronic structure of the ligand. This representation captures the essential chemical properties that influence binding interactions, such as hydrogen bonding capacity, hydrophobic character, and electrostatic potential.

The virtual joint node serves as a computational hub that connects all protein residues and all ligand atoms, creating a bridge between the two molecular entities. This node acts as an information exchange center, allowing the model to learn the complex patterns of protein-ligand interactions without requiring explicit pairwise connections between every residue and atom. The virtual joint node can be interpreted as representing the binding pocket itself, a spatial region where protein-ligand interactions occur, providing a biologically meaningful representation of the binding interface.

### Node Feature Encoding

The input graph *G* = (*V, E*) consists of three distinct node types: protein residues, ligand atoms, and a special joint node that serves as a bridge between the protein and ligand domains. Each node *v*_*i*_ ∈ *V* is characterized by a feature vector 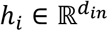, where *d*_*in*_ = *N*_*residue*_ + *N*_*atom*_ + 1 represents the concatenated one-hot encodings of residue types, atom types, and the joint node indicator.

The node features are constructed as follows:

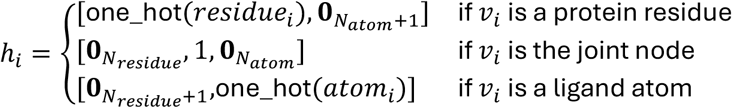

### Edge Feature Encoding

The edge set *E* encompasses multiple interaction types that capture the complex spatial and chemical relationships within the protein-ligand system. For each edge *e*_*ij*_ ∈ *E*, the edge features 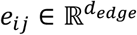 encode distance information, structural connectivity, and chemical bond properties:

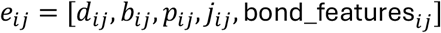

where *d*_*ij*_ represents the Euclidean distance between nodes *i* and *j, b*_*ij*_ is a binary indicator for backbone connectivity, *p*_*ij*_ indicates protein-joint connections, *j*_*ij*_ represents joint-ligand connections, and bond_features_*ij*_ contains the one-hot encoding of chemical bond types for ligand atoms.

### Spatial Contact Detection

We employ a grid-based spatial contact detection algorithm to efficiently identify protein residues within a distance threshold *τ*_*contact*_. For each residue pair (*i, j*), if their Cα atoms satisfy ||x_*i*_ − x_*j*_||_2_ < *τ*_*contact*_, an edge is established. This approach reduces computational complexity from *O*(*n*^2^) to *O*(*n*) through spatial partitioning, enabling the model to handle large protein structures efficiently while maintaining the essential spatial relationships that govern protein-ligand interactions.

### Graph Convolutional Layer Architecture

The core of YuelPocket consists of multiple Graph Convolutional Layers^34^ (GCL) that iteratively update node and edge representations. Each GCL layer implements a message-passing mechanism where node features are updated through the aggregation of messages from neighboring nodes. The edge model computes updated edge features:

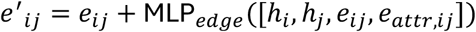

where MLP_*edge*_ is a multi-layer perceptron with LayerNorm and SiLU activation functions. The node model aggregates messages from connected edges and updates node representations:

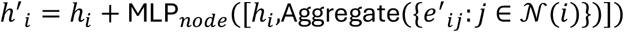

The aggregation function employs either sum or mean pooling with normalization:

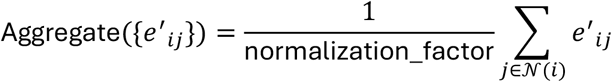

### Multi-Layer Architecture and Output Prediction

The complete GNN architecture stacks *L* GCL layers with residual connections, where each layer maintains the hidden dimension *d*_*hidden*_. The final node representations are projected to output dimensions through linear transformations:

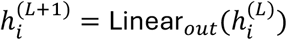

For pocket prediction, the model outputs a probability score for each protein residue:

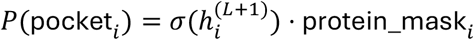

where *σ* is the sigmoid activation function and protein_mask_*i*_ ensures that predictions are only made for protein residues. The model is trained using binary cross-entropy loss:

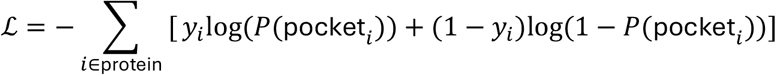

where *y*_*i*_ represents the ground truth pocket labels for each residue.

### Pocket Center Detection Algorithm

The pocket center detection algorithm employs a sophisticated FFT-based docking-like scoring approach that combines protein structure information with machine learning predictions to identify potential binding sites. The algorithm operates on a three-dimensional grid representation of the protein structure, where each grid point is evaluated for its suitability as a pocket center based on both geometric constraints and predicted pocket probabilities.

The algorithm begins by constructing a 3D grid around the protein structure with a configurable spacing (default 0.5 Å) and padding (15 Å) to ensure comprehensive coverage of potential binding sites. Each protein atom is then mapped to the grid, with atoms assigned different scoring values based on their distance from the atom center and the associated residue’s pocket prediction score. Specifically, atoms create a dual-zone scoring system: an inner repulsion zone (0 to 1.5 Å) marked with negative values (−100.0) to prevent pocket centers from being placed too close to protein atoms, and an outer attraction zone (1.5 to 2.0 Å) where grid points receive positive scores based on the pocket prediction probabilities of the corresponding residues.

The core innovation of this algorithm lies in its use of FFT convolution to efficiently evaluate potential pocket locations. Two probe kernels are created: a primary kernel with a radius of 2.0 Å for identifying attractive regions, and a secondary kernel with a radius of 5.0 Å for broader spatial analysis. The FFT convolution process allows the algorithm to rapidly compute scoring grids by transforming the protein grid and probe kernels to the frequency domain, performing element-wise multiplication, and transforming back to the spatial domain. This approach significantly reduces computational complexity compared to traditional distance-based calculations.

Following the FFT scoring, the algorithm identifies grid points with positive scores as potential pocket locations and applies a hierarchical clustering approach using DBSCAN^31^ (Density-Based Spatial Clustering of Applications with Noise). The clustering process groups nearby high-scoring points into coherent pocket regions, with parameters optimized for biological relevance (eps = 2.5 Å, min_samples = 30). For large clusters containing more than 200 points, the algorithm implements a secondary clustering step that refines the pocket centers by selecting the top 30 points based on secondary scores and performing additional DBSCAN clustering with relaxed parameters (eps = 4 Å, min_samples = 1). The final pocket centers are determined as the spatial centroids of the resulting clusters, with associated confidence scores calculated as the mean scores of all points within each cluster.

This multi-stage approach ensures that the detected pocket centers represent biologically meaningful binding sites rather than isolated high-scoring grid points, while the FFT-based scoring provides computational efficiency necessary for large-scale protein analysis. The algorithm’s ability to integrate machine learning predictions with structural constraints makes it particularly effective for identifying both canonical binding sites and novel allosteric pockets that may not be immediately apparent from structural analysis alone.

### Probe Selection Algorithm

The probe selection algorithm is designed to efficiently identify a minimal set of ligands (probes) that collectively maximize the coverage of true binding pocket residues across a diverse set of protein targets. The process operates in a batch-wise manner, iteratively selecting ligands that contribute the most new coverage of pocket residues for each batch of proteins. For each batch, the algorithm first determines the set of true pocket residues for the proteins in the batch. It then initializes the current coverage using predictions from the already selected probe set. If the current coverage already exceeds a predefined threshold (e.g., 80 %), the batch is skipped. Otherwise, the algorithm evaluates each candidate ligand by predicting its pocket coverage across the batch and quantifies how many new, previously uncovered, true pocket residues it identifies. Ligands that contribute new coverage are added to the probe set, and the process continues until the coverage threshold is met or all ligands are exhausted. This greedy, coverage-driven approach ensures that the selected probe set is both efficient and effective in representing the diversity of binding pockets in the dataset.

### Metrics

We evaluate YuelPocket’s performance using standard classification metrics, including ROC curves (AUC-ROC), precision-recall curves (AUC-PR), precision 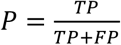,recall 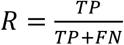, specificity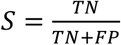, negative predictive value 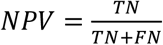, Matthews correlation coefficient 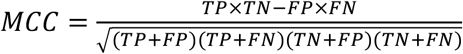, and F1-score 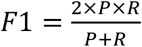. For comparison with traditional pocket prediction methods, we use distance center-to-center *DCC* = ||**x**_*pred*_ − **x**_*ligand*_||_2_ and success rates considering top N and N+2 ranked pocket centers, where N represents the number of true binding sites per protein. To assess correlations between pocket probabilities and binding affinities, we employ Pearson correlation and Spearman correlation coefficients with p-values for statistical significance testing.

## ACKNOWLEDGMENTS

We acknowledge support from the National Institutes of Health 1R35 GM134864 and the National Science Foundation grant 2210963.

## DATA AND SOFTWARE AVAILABILITY

Source codes and test data are deposited at: https://github.com/hust220/yuel_pocket.git.

## SUPPORTING INFORMATION AVAILABLE

The supporting information provides Figure S1-S3, and Tables S1-S4.

## AUTHOR CONTRIBUTIONS

Jian Wang contributed to the conceptualization, methodology, model development, data analysis, writing of the original draft, and visualization of the study. Nikolay V. Dokholyan provided supervision, resources, writing review and editing, project administration, and funding acquisition.

