## Supplemental Information for "Unified Protein-Small Molecule Graph Neural Networks for Binding Site Prediction"

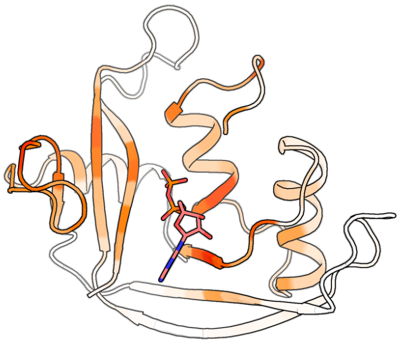

**Figure S1. Pocket residues of PDB 1HI5 predicted by YuelPocket.**

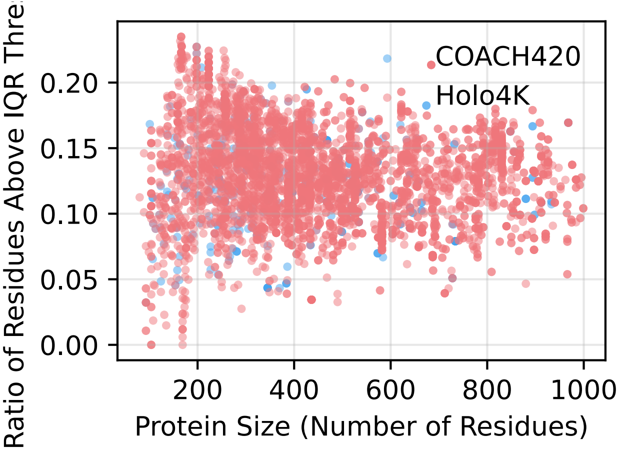


**Figure S2. Ratio of residues above adaptive-threshold versus protein size.**

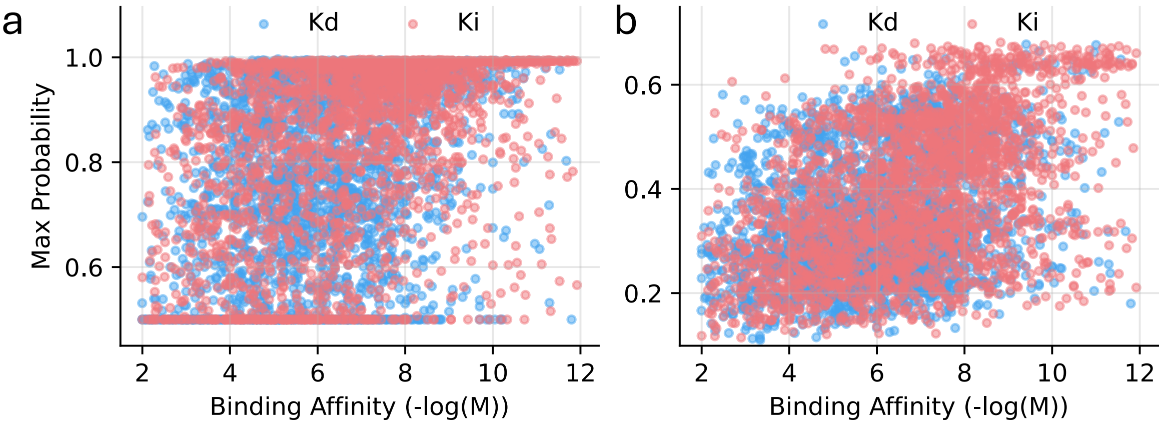


**Figure S3. Correlation between pocket probability and binding affinity.**

**Table S1. Evaluation of distance "center–center" (DCC), top N, top N + 2 scores of YuelPocket.**

| Distance Cutoff (Å) | DCC | N+2 | N |
|---|---|---|---|
| 0.000 | 0.000 | 0.000 | 0.000 |
| 0.500 | 0.156 | 0.121 | 0.066 |
| 1.000 | 0.495 | 0.409 | 0.238 |
| 1.500 | 0.610 | 0.516 | 0.309 |
| 2.000 | 0.649 | 0.555 | 0.336 |
| 2.500 | 0.681 | 0.587 | 0.358 |
| 3.000 | 0.707 | 0.610 | 0.372 |
| 3.500 | 0.734 | 0.633 | 0.390 |
| 4.000 | 0.760 | 0.660 | 0.406 |
| 4.500 | 0.773 | 0.671 | 0.420 |
| 5.000 | 0.787 | 0.689 | 0.432 |
| 5.500 | 0.801 | 0.708 | 0.447 |
| 6.000 | 0.822 | 0.730 | 0.468 |
| 6.500 | 0.837 | 0.755 | 0.501 |
| 7.000 | 0.854 | 0.774 | 0.524 |
| 7.500 | 0.868 | 0.793 | 0.546 |
| 8.000 | 0.883 | 0.810 | 0.569 |
| 8.500 | 0.895 | 0.831 | 0.593 |
| 9.000 | 0.907 | 0.846 | 0.612 |
| 9.500 | 0.920 | 0.861 | 0.630 |
| 10.000 | 0.930 | 0.877 | 0.656 |
| 10.500 | 0.936 | 0.889 | 0.681 |
| 11.000 | 0.942 | 0.901 | 0.704 |
| 11.500 | 0.948 | 0.911 | 0.729 |
| 12.000 | 0.951 | 0.919 | 0.746 |
| 12.500 | 0.956 | 0.927 | 0.761 |
| 13.000 | 0.961 | 0.933 | 0.779 |
| 13.500 | 0.965 | 0.939 | 0.797 |
| 14.000 | 0.965 | 0.942 | 0.813 |
| 14.500 | 0.969 | 0.948 | 0.828 |
| 15.000 | 0.971 | 0.951 | 0.844 |
| 15.500 | 0.973 | 0.954 | 0.855 |
| 16.000 | 0.974 | 0.956 | 0.864 |
| 16.500 | 0.976 | 0.959 | 0.874 |
| 17.000 | 0.977 | 0.961 | 0.884 |
| 17.500 | 0.979 | 0.965 | 0.893 |
| 18.000 | 0.980 | 0.967 | 0.902 |
| 18.500 | 0.980 | 0.968 | 0.909 |
| 19.000 | 0.980 | 0.969 | 0.915 |

| | | | |
|---|---|---|---|
| 19.500 | 0.982 | 0.971 | 0.918 |
| 20.000 | 0.983 | 0.974 | 0.925 |

**Table S2. Comparison of distance "center–center" (DCC), top N, top N + 2 scores among YuelPocket, SiteRadar, Fpocket, and PUResNet.**

| | DCC | N+2 | N |
|---|---|---|---|
| YuelPocket | 0.787 | 0.689 | 0.432 |
| SiteRadar | 0.760 | 0.720 | 0.490 |
| Fpocket | 0.710 | 0.460 | 0.310 |
| PUResNet | 0.460 | N/A | 0.470 |

**Table S3. The 15 probes in the minimal probe set.**

| PDB_ID | SMILES | size |
|---|---|---|
| 1nh0_0 | OC[C@H](O)[C@H](CC1CCCCC1)N[C@H](O)OCC1CCCCC1 | 23 |
| 1ofz_2 | C[C@@H]1O[C@H](O)[C@@H](O)[C@H](O)[C@@H]1O | 11 |
| 2bxm_0 | CCCCCCCCCCCCCC(O)O | 16 |
| 3ar4_2 | CC(O)OC[C@H](CO[PH](O)(O)OCCN)OC(C)O | 19 |
| 3cbg_1 | COC1CC(CCC(O)O)CCC1O | 14 |
| 3evd_0 | NC1NC(O)C2NCN([C@@H]3O[C@H](CO[PH](O)(O)O[PH](O)(O)O [PH](O)(O)O)[C@@H](O)[C@H]3O)C2N1 | 32 |
| 3fp0_1 | CC1CC[C@@]2(CO)[C@@H](C1)O[C@@H]1[C@H](OC(C)O) C[C@@]2(C)[C@]12CO2 | 22 |
| 3k5i_2 | NC1CNCN1[C@@H]1O[C@H](CO[PH](O)(O)O)[C@@H](O)[C@H]1O | 19 |
| 3k5i_4 | N[C@H]1CNCN1[C@@H]1O[C@H](CO[PH](O)(O)O)[C@@H](O)[C@H ]1O | 19 |

| | | |
|---|---|---|
| 3nr4_1 | OS(O)(NCC1CCCCN1)C1CCC(Br)C2CCCCC21 | 22 |
| 4csj_0 | CC1C[C@@H](C)C[C@H](C)C1S(O)(O)N[C@@H](C)CNC1CCCC2C1CNN2C1CCC(F)CC1 | 33 |
| 4rf9_0 | NC(N)NCCC[C@H](N)C(O)O | 12 |
| 4yha_1 | CC(O)NC1S[C@@H](S(N)(O)O)NN1C | 14 |
| 5tvf_1 | NC(NNCC1CCCC(C(N)N)C1)NNCC1CCCC(C(N)N)C1 | 26 |
| 6fu1_0 | CCCCCC(O)N(O)[C@H]1C[C@H](O)N(C2CCC(Cl)CC2)C1O | 23 |

**Table S4. Correlation between pocket probability and the binding affinity in the PDBBind dataset.**

| Analysis Method | Affinity Measure | Sample Size | Pearson Correlation | Spearman Correlation |
|---|---|---|---|---|
| Maximum Probability | All Samples | 5314 | 0.391 (p=1.444e-193) | 0.423 (p=1.596e-229) |
| Maximum Probability | Kd | 2783 | 0.302 (p=1.383e-59) | 0.309 (p=1.563e-62) |
| Maximum Probability | Ki | 2531 | 0.428 (p=4.163e-113) | 0.469 (p=1.173e-138) |
| Mean Probability | All Samples | 5314 | 0.429 (p=4.013e-237) | 0.415 (p=8.881e-220) |
| Mean Probability | Kd | 2783 | 0.306 (p=1.996e-61) | 0.290 (p=6.393e-55) |
| Mean Probability | Ki | 2531 | 0.476 (p=2.310e-143) | 0.470 (p=1.340e-139) |
